# Reconstructing phylogeny from reduced-representation genome sequencing data without assembly or alignment

**DOI:** 10.1101/225623

**Authors:** Huan Fan, Anthony R. Ives, Yann Surget-Groba

## Abstract

Although genome sequencing is becoming cheaper and faster, reducing the quantity of data by only sequencing part of the genome lowers both sequencing costs and computational burdens. One popular genome-reduction approach is restriction site associated DNA sequencing, or RADseq. RADseq was initially designed for studying genetic variation across genomes usually at the population level, and it has also proved to be suitable for interspecific phylogeny reconstruction. RADseq data pose challenges for standard phylogenomic methods, however, due to incomplete coverage of the genome and large amounts of missing data. Alignment-free methods are both efficient and accurate for phylogenetic reconstructions with whole genomes and are especially practical for non-model organisms; nonetheless, alignment-free methods have only been applied with whole genome sequences. Here, we test a full-genome assembly and alignment-free method, AAF, in application to RADseq data and propose two procedures for reads selection to remove missing data. We validate these methods using both simulations and a real dataset. Reads selection improved the accuracy of phylogenetic construction in every simulated scenario and the real dataset, making AAF comparable to or better than alignment-based method with much lower computation burdens. We also investigated the sources of missing data in RADseq and their effects on phylogeny reconstruction using AAF. The AAF pipeline modified for RADseq data, phyloRAD, is available on github (https://github.com/fanhuan/phyloRAD).

## Introduction

With the introduction of Next-Generation Sequencing (NGS) technologies, phylogenies are being increasingly reconstructed using genomic data rather than a few loci. Genomic data provide more information about the evolutionary history of the study group, although this comes with the costs of higher sequencing expenses and heavier computational burdens. For ecological or evolutionary studies, especially at the population level, it is still expensive to sequence full genomes of tens or hundreds of individuals, and it is difficult to process such large quantities of data. Therefore, there is much current interest in methods that target only part of the genomes. By removing unwanted information such as repetitive elements (Barbazuk *et al*. 2005) and focusing on certain regions, these reduced-representation approaches allow the pooling of many samples in one sequencing lane while still achieving high sequencing coverage (Arnold *et al*. 2013).

The most common type of reduced representation approaches uses restriction site associated DNA (RAD). Despite variation in library preparation (reviewed in Davey *et al*. 2011; Andrews *et al*. 2016), these methods only sequence the flanking regions of restriction sites. Since no prior genetic or genomic information is needed, these methods are popular for rapid genetic marker discovery and genotyping for non-model organisms that usually lack reference genomes. Although not the original intention, RADseq data have been used for phylogeny reconstructions, both among and within species (Rubin *et al*. 2012; Cruaud *et al*. 2014; Emerson *et al*. 2010).

Despite the reduction in data quantity, analysis of RADseq data is not much easier than whole-genome data. Two of the biggest challenges are (i) clustering reads from the same restriction site quickly and accurately in the presence of both polymorphism and sequencing errors, and (ii) de-novo assembling each cluster into unique loci despite the existence of paralogues and repetitive sequences (Chong *et al*. 2012). The difficulties of clustering and assembly are similar to the difficulties of alignment and assembly in standard whole-genome analyses. Therefore, assembly and alignment-free methods that are efficient in phylogeny reconstruction from whole genome datasets (Sims *et al*. 2009; Yi & Jin 2013; Fan *et al*. 2015) could have advantages for analyzing RADseq data. These methods use short strands of nucleotides or amino acids, usually referred to as k-mers (*k* being the length of the DNA or protein strand). Because the methods are based on fragments of the genome, there is no need for assembly and alignment, which is a particular advantage for non-model organisms due to their lack of reference genomes. The method described in Fan et al. (2015), “AAF”, is designed for phylogenomic analysis of large eukaryote genomes, which is the arena for RADseq data. The ability of AAF to reconstruct phylogenies without the need either to cluster reads or to assemble clusters makes it a natural candidate for analyzing RADseq data.

Due to the differences in library construction (Lepais & Weir 2014), such as the number of restriction endonuclease applied, whether or not to apply restriction exclusion (RESTseq, Stolle & Moritz 2013) and size selection, there are a number of different types of RADseq data. While sequencing errors and uneven coverage are problems shared with shotgun sequencing of whole genomes using NGS technologies, an extra challenge of phylogenetic reconstruction using any RADseq technology is missing data caused when information from a restriction site is missing in one or more samples in the final data matrix (Arnold *et al*. 2013). Information from these restriction sites could dropout in two ways. First, when there is a mutation within a restriction site, its flanking regions will not be sequenced due to the absence of digestion. This could also affect the neighboring restriction sites if there is a size selection step downstream (Figure 2a in Andrews *et al*. 2016). This type of dropout is not random since it follows evolutionary history, and it has been previously well investigated (Arnold *et al*. 2013). There could also be random dropout in library construction due to random shearing, methylation or UV light exposure (Mastretta-Yanes et al. 2015), or in the bioinformatic analysis due to uneven coverage (Catchen *et al*. 2013). Random dropouts, however, have not been systematically investigated as a source of errors in phylogenetic reconstruction using RADseq data. Missing data always present problems for phylogeny reconstruction, but the problems are much larger for RADseq data (reviewed in Andrews et al. 2016), and the distribution of missing data across samples varies significantly among loci (Figure 1 in Huang & Knowles 2016). Therefore, a common practice is to exclude loci that have data missing from more than a threshold number of samples (e.g. Zellmer *et al*. 2012). The higher the threshold, the more conservative is the approach, ensuring greater representation of loci across samples at the expense of reducing information shared by some but not all samples. However, because nonrandom and random loss of information are hard to separate, they can lead to unforeseen consequences whether they are excluded or not (Huang & Knowles 2016).

**Figure 1.**
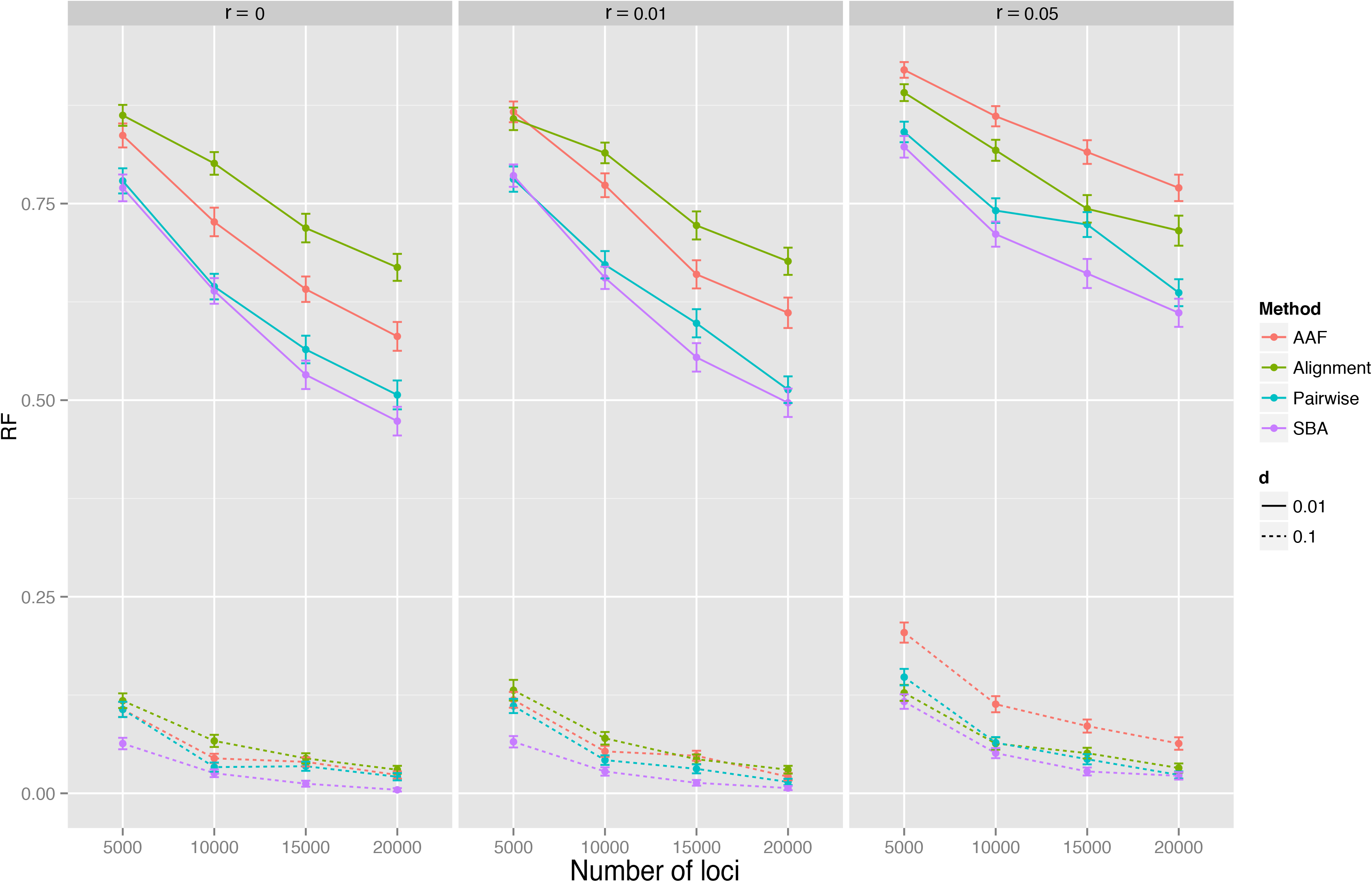
Normalized RF distances between phylogenetic trees reconstructed using different numbers of RAD loci and the 12-sample starting phylogeny. Results for RADseq data with a deeper starting phylogeny (*d = 0.1*) are denoted by solid lines, and those for a shallower starting phylogeny (*d = 0.01*) are denoted by dashed lines. Line colors denote different methods, and the three vertical panels are for different random dropout rates. Each data point is the mean of 100 simulations.

**Figure 2.**
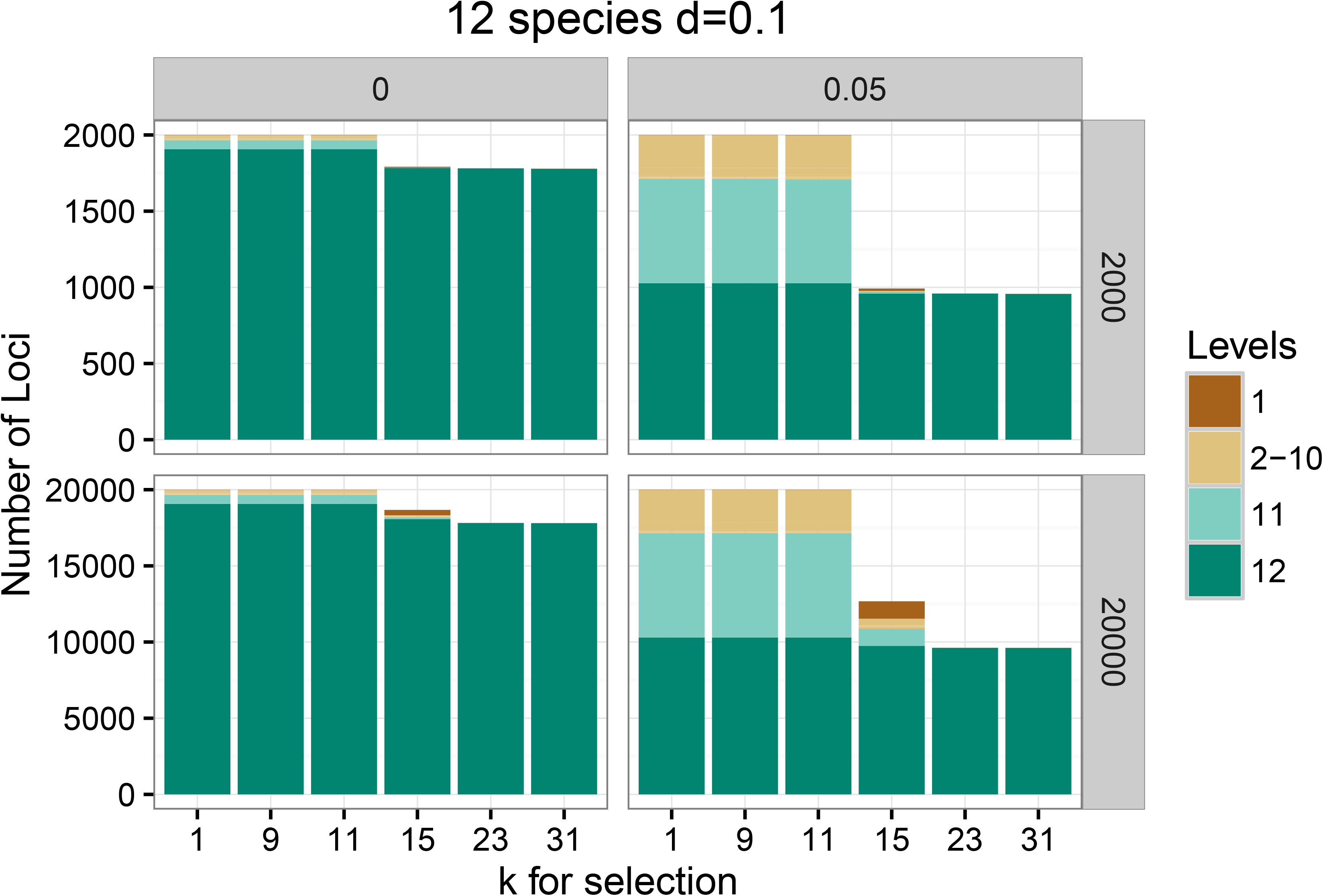
Composition of different type of RAD loci as a function of *k_s_* used in SBA reads selection with or without random dropout for datasets of different sizes. The simulations were based on the deep 12-species starting phylogeny (d = 0.1). Different colors represent different sample counts at each locus. For example, 12 (the dominant pink color) represent the RAD loci that are shared by all the 12 samples. The vertical axis gives the number of RAD loci in each count level, and the horizontal axis gives the *k_s_* used for SBA reads selection. Note that all loci appearing in fewer than 12 samples would not be included in SBA reads selection, but they could be included in pairwise reads selection; for example, a locus that occurs in 2 samples would be included in the pairwise calculation of the distance between them.

In this study, we test the applicability of AAF to RADseq data. We also develop and validate a reads-selection step that is added to the original AAF pipeline to address the missing data in RADseq. This step assumes that reads from different samples came from the same restriction-site containing locus, or RAD locus, if they contain at least one (singleend sequences) or two (paired-end sequences) *k*-mers in common. Our first type of reads selection only picks reads from RAD loci that are identified as being shared by all (SBA) samples. This is done by first identifying *k*-mers that are shared by all the samples, and then keeping reads in each sample that contain those *k*-mers. For large sample sizes or large volumes of missing data where the number of SBA loci is limited, we develop a second method that computes distances between pairs of samples after first selecting reads based on shared *k*-mers between this pair. This pairwise selection retains more information than SBA, because the information loss is independent of the total number of samples. We demonstrate how these two types of reads selection improve the performance of the original AAF method using both simulations and a real dataset of 34 individuals from six *Quercus* species (Cavender-Bares *et al*. 2015).

## Methods and materials

### AAF and reads selection

AAF (Fan *et al*. 2015) computes pairwise distances between samples from the numbers of shared and total k-mers using an evolutionary model of mutation (including substitutions, insertions, and deletions). The k-mers used in AAF are overlapping strands of DNA that are generated from window-sliding down the reads with a window size of *k*; for example, a read of length 10 would generate six 5-mers. The AAF pipeline consists of four steps: (i) identifying all the k-mers from each sample, (ii) calculating the number of shared *k*-mers between each pair of samples, (iii) calculating the distances between each pair using the number of shared and total *k*-mers, and (iv) reconstructing the phylogeny from the distance matrix using a modified Fitch-Margoliash method (Fitch and Margoliash, 1967).

When applied to RADseq data, RAD loci dropouts will be interpreted as large deletions, since the flanking regions around these restriction sites will not be sequenced. While nonrandom dropout will create deletions that contain phylogenetic information, random dropout will create deletions that are independent of the phylogeny and are therefore not included in the evolutionary model on which AAF is based. Because it is not easy to separate the two types of dropout in real datasets, we take a conservative approach to account for dropouts in RADseq data using two types of reads selection.

### Reads selection: Shared By All (SBA)

The first approach for dealing with missing data selects reads that contain *k*-mers in common across all samples. If the dataset is single-ended, reads that contain at least one k-mer that are SBA are kept. If the dataset is pair-ended, a pair is kept if both reads contain at least one SBA k-mer. By increasing the length of k-mers used for selection (denoted *k_s_*), selection is made stricter. The performance of the method depends on *k_s_*: *k_s_* has to be large enough to exclude homoplasy in which reads from different RAD loci are treated as being from the same locus, and *k_s_* must be small enough so that reads from the same RAD locus share at least one *k*-mer without mutations.

Datasets containing many taxa and high dropout rates may not be appropriate for SBA reads selection, because the combination of nonrandom and random dropout will leave only a few SBA loci. For example, a dropout rate of 0.05 would reduce the amount of genetic information in a 10-sample data set to 60% (=[1 – 0.05]^10^) and a 100-sample data set to <1% (=[1 – 0.05]^100^).

### Reads selection: Pairwise

The second type of reads selection is similar to SBA but selects reads on a pairwise basis. This is possible because AAF uses a distance method to construct phylogenies, so pairwise reads selection can be performed before pairwise distance computation. This reads selection is conducted the same way as SBA reads selection except that *k*-mers used are shared by each pair instead of all the samples.

### Simulation of RADseq data

To test AAF and reads selection, we simulated RADseq sequences using a coalescent-based RADseq simulator called *simrrls* (https://github.com/dereneaton/simrrls). We adopted the default tree topology from the program, which is a 12-taxon balanced tree, and to assess different levels of divergence, we scaled the phylogeny to have an average root to tip length of either 0.1 (d = 0.1) or 0.01. The coverage follows a normal distribution with a mean of 8 and a standard deviation of 4; these settings lead to a 2% chance of a restriction site locus having no coverage (missing), and a 4% chance of having no coverage or single coverage. This coverage level allows AAF (after reads selection) to filter k-mers, that is, to include *k*-mers in a given sample only if they appear at least twice (Fan *et al*. 2015); this filtering effectively reduces data contamination caused by sequencing error. Note that all the AAF analysis done in this study is with *k*-mer filtering, since higher coverage is a feature of RADseq data. Sequencing error was set to match current Illumina platforms (0.1%, NGS field guide, 2014), and other parameters are set as defaults: read length (100 base pairs including 6 base pairs of barcode), per-site mutation rate (1e-9), and effective population size (5e5). The number of RAD loci that is simulated sets the size of the dataset.

*simrrls* simulates both loss and gain of restriction sites due to mutations. Mutations within existing restriction sites result in the loss of flanking regions in the RADseq data (option −mc). Mutations can give rise to restriction sites, and consequently their flanking regions will be included in the RADseq data (option −ms). Random dropout of loci due to library construction is not included in *simrrls*, so we added it in the downstream processing using a custom python script. The random dropout rates we used in the simulation study were *r* = 0.01 and 0.05 (meaning that for each individual, 1% and 5% of the loci will be removed randomly from the simulated dataset). Random dropout rate has not been studied systematically. Nonetheless, Baird et al. (2008) generated RADseq data from 41,622 loci from three pooled populations using restriction enzyme *SbfI*, which was lower than the 44,709 restriction sites that exist in the reference genome. These results imply that 7% of loci are missing due to a combination of polymorphism in restriction sites among individuals (nonrandom dropout) and random dropout. This was one of the earliest RADseq studies, and therefore we consider this to be at the extreme high end of random dropouts. Therefore, we selected 1% and 5% to represent low and high random dropout rates for our simulations.

Each scenario for a specified combination of parameters was simulated 100 times.

### Alignment method and tree comparisons

In simulations, the accuracy of AAF was measured on both absolute and relative scales by comparing the AAF phylogenies to the starting phylogeny used to simulate the RADseq data, and to phylogenies constructed using the aligned data. For aligning, the data were concatenated to give the alignment at each simulated locus. *simrrls* simulates reads of fixed length on a per-locus basis and keeps track of locus information for each read. Because we did not simulate indels, it was straightforward to generate an alignment for each locus. When there were heterozygous positions within a sample, we coded it as ambiguous (Emerson *et al*. 2010) with a threshold of 70%: if the majority state appears in less than 70% of the reads containing that locus for an individual sample, then that position was coded as X in the consensus. Missing loci in each sample were represented by gaps. After obtaining the total alignment, pairwise distances between samples were calculated with the F81 model in dnadist from the PHYLIP package (v 3.69, Felsenstein 2005). Then the phylogeny was reconstructed using the program fitch in PHYLIP; this is similar to the distance method as used in the AAF method. The accuracy of the phylogenies from each method was measured using the normalized Robinson-Foulds (RF) distance (Robinson & Foulds 1981; Bernard *et al*. 2016), with 0 indicating an identical phylogeny to the starting phylogeny used for the simulations, and 1 being random.

### Real dataset

We validated the methods with two real RADseq datasets. The first one consists of 34 individuals from six *Quercus* species, presenting a mixture of inter- and intra-specific relationships. This dataset was originally published by Cavender-Bares *et al*. (2015) under the NCBI bioproject PRJNA277574. Details about this dataset can be found in Table S1. The second datasets consists of more samples and a deeper divergence, with 74 individuals from 17 genus of the Phrynosomatidae subfamily. This dataset is obtained from Leache *et al*. (2015), and details about it can be found in Table S2.

## Results

### AAF works as well as the alignment-based method on RADseq data

When there was no or low (*r* ≤ 0.01) random dropout of RAD loci, AAF slightly outperformed the alignment method with both deep (*d* = 0.1) and shallow (*d* = 0.01) phylogenies (i.e., lower RF distances between the reconstructed phylogeny and the starting phylogeny in the simulations, Figure 1). When the random dropout rate was high (*r* = 0.05), the alignment method performed slightly better than AAF.

Increasing the number of RAD loci yielded better results in general (Figure 1), as is the case for AAF applied to genome data (Fan et al. 2015, Figure 3). Furthermore, when the starting tree used for simulations was deep (d = 0.1), all methods performed well even with the smallest dataset tested (5000 loci = 500k base pairs), and their resulting phylogenies were close to the starting phylogeny with 20,000 loci. On the other hand, the RF distances between reconstructed phylogenies and the starting phylogeny were much higher with the shallower starting phylogeny (d = 0.01), and 20,000 RAD loci did not provide enough information for accurate phylogeny reconstruction. This suggests more RAD loci are needed to reconstruct shallow phylogenies, which is intuitive since less phylogenetic information will be available for recently diverged taxa.

**Figure 3.**
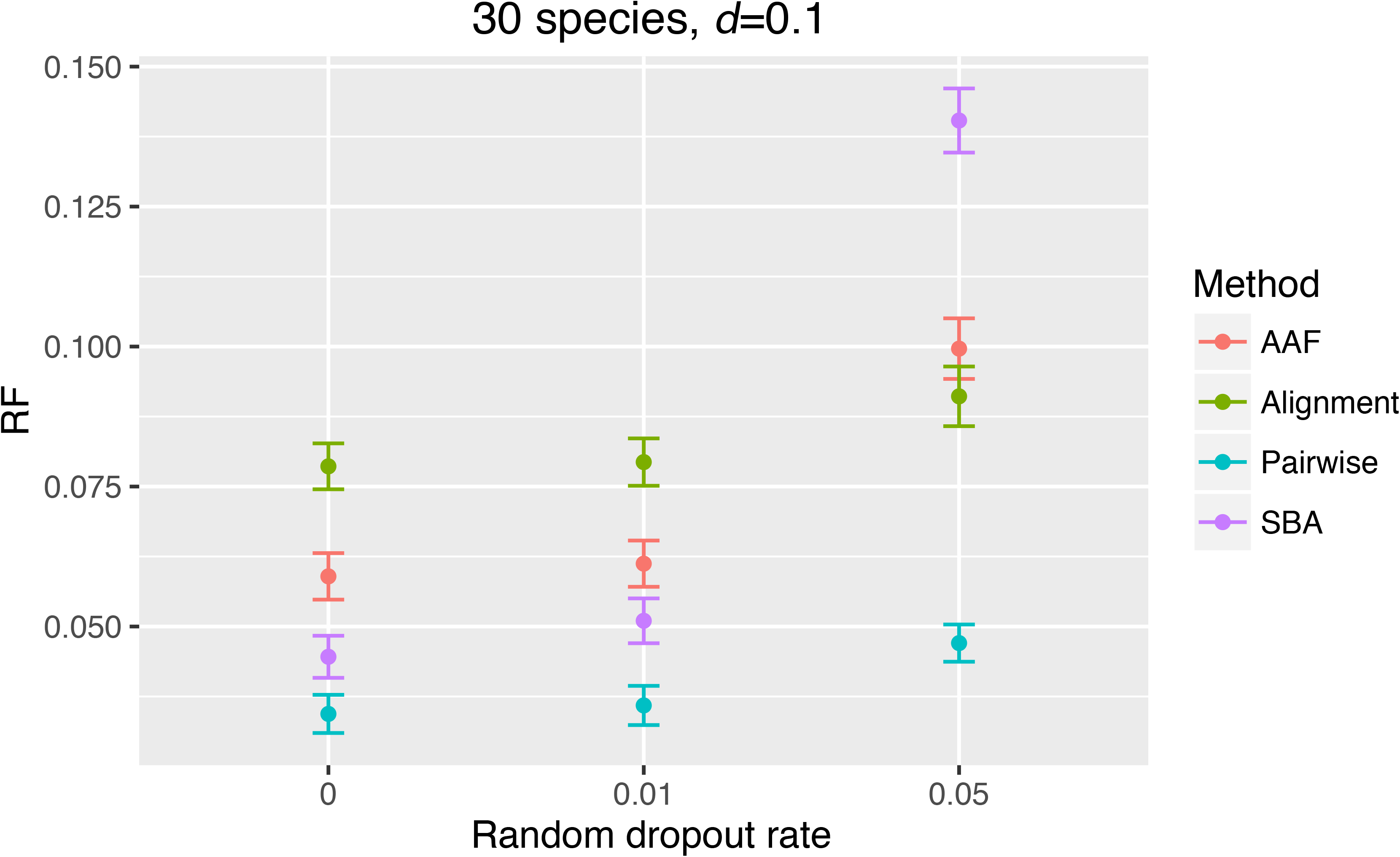
Normalized RF distance between phylogenetic reconstructions and the phylogeny used to simulate the datasets using different *k_s_* and a 30-tip random starting phylogeny with a root-to-tip average depth of *d =* 0.1. One hundred simulations were performed for each scenario.

### Reads selection improves the performance of AAF

Reads selection proved to be a very effective way to eliminate missing data when *k_s_* is large enough (Figure 2), and this pre-processing improved the performance of AAF, either with SBA or pairwise (Figure 1). When the dropout rate was high (*r* = 0.05), reads selection was required for AAF to outperform the alignment method. Reads selection also helped the performance of AAF in the absence of random dropouts (*r* = 0).

When the number of samples was low (12, Figure 1), SBA performed slightly better than pairwise reads selection. When the number of samples was higher (30, Figure 3), pairwise reads selection performed better, especially when the random dropout rate was high.

### *k* for selection (*k_s_*) and for phylogeny reconstruction (*k_p_*)

The effect of reads selection can be assessed from the coverage of samples at each RAD locus after reads have been selected (Figure 2); the simulation was done on a locus-basis therefore it was possible to keep track of the loci from which reads originate. AAF with no reads selection corresponds to *k_s_* = 1, because every read at least contains one of the four nucleotides. Reads selection with *k_s_* = 9 and 11 had little effect on the number of samples sharing reads from the same RAD loci (Figure 2) because of homoplasy: a read will be retained in the dataset if it contains a *9*-mer or 11-mer that is found in reads from the other 11 samples regardless of whether these reads come from the same RAD locus. As a result, there is no improvement in phylogenetic reconstructions compared to no reads selection (Figure 4). For the smaller dataset (2,000 RAD loci), *k_s_* = 15 greatly reduced the number of reads that were not shared by all samples (Figure 2), resulting in better performance in phylogenetic reconstruction (Figure 4). For the larger dataset (20,000 loci), *k_s_* = 15 removed many of the reads that were not shared by all RAD loci, although for the case of high random dropout, many reads remained that were unique to single samples due to homoplasy. This resulted in no better, if not worse, performance in reconstructing the phylogeny (Figure 4). However, *k_s_* = 23 was long enough to overcome homoplasy, leading to all selected loci being those that are in fact shared by all samples. This led to peak performance for phylogenetic reconstruction. Larger *k_s_* (*k_s_* = 31) also overcame homoplasy and led to all loci being shared by all, yet yielded fewer reads (Figure S1). This means not all SBA reads were kept. This might due to more than one mutation within *k* base pairs, and this is more likely when *k* is longer. The performances of phylogenetic reconstruction are consequentially slightly reduced in some scenarios.

**Figure 4.**
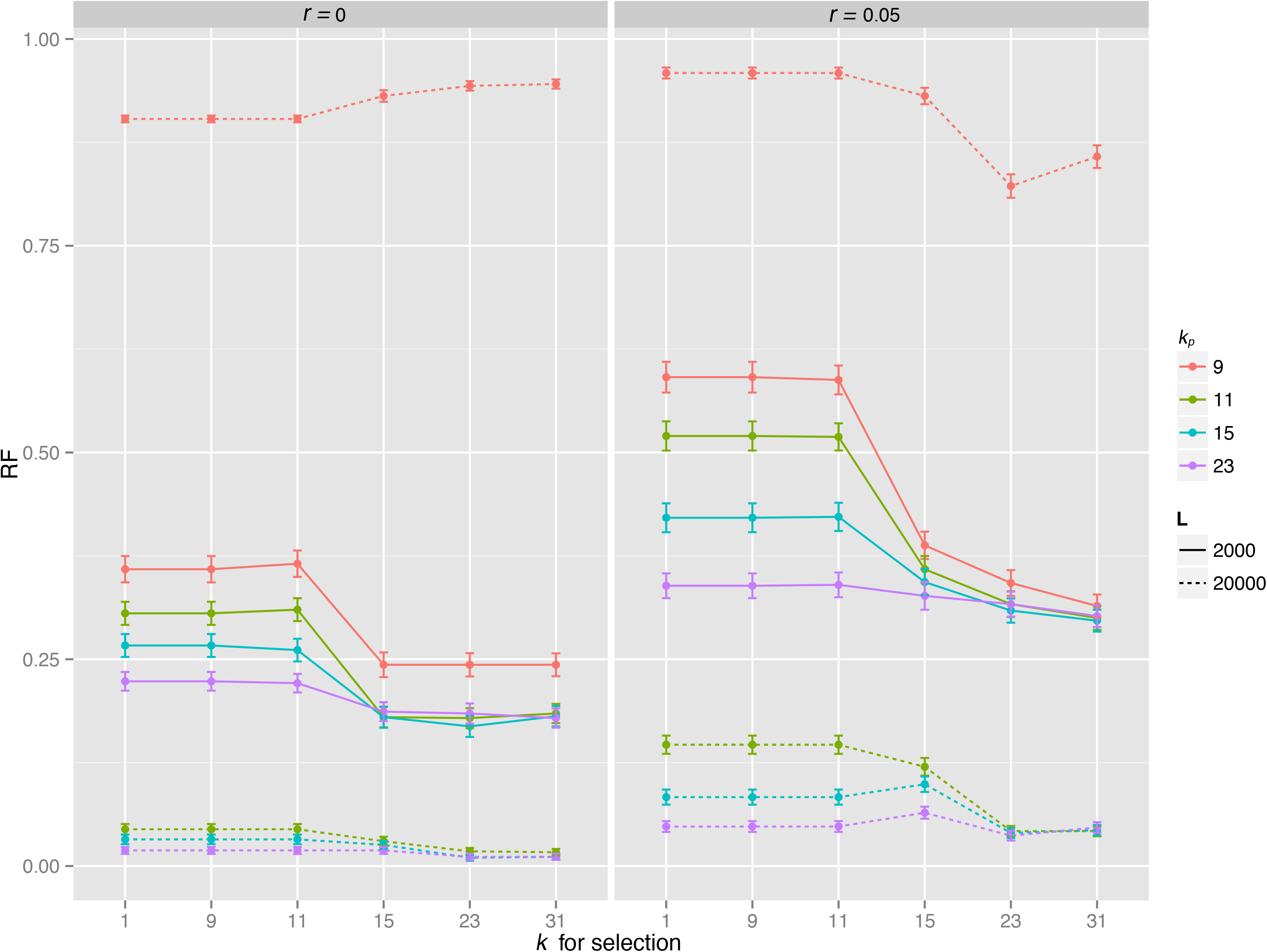
NormalizedRF distance between phylogenetic reconstructions and the phylogeny used to simulate the datasets with different combinations of *k_p_* and k_s_. Note that *k_s_=1* represents AAF without reads selection. The simulations are based on a starting phylogeny of 12 tips with an average root to tip length of *d =* 0.1, with either *L =* 2000 or 20000 loci, and a random dropout rate of *r =* 0 or 0.05. One hundred simulations were performed for each scenario.

When there is ineffective or no reads selection, larger *k_p_* for construction the phylogeny from the reads yielded better results, because larger *k_p_* guards against homoplasy at the reconstruction phase. For *k_s_* large enough to have effective reads selection (*k_s_* = 23 or 15 for small and large datasets), the performance of reconstruction became stable once *k_p_* was over the optimal value estimated for AAF when applied to whole genomes (Figure 2a in Fan *et al*., 2015).

### Random dropout

The increase in the random dropout rate resulted in an increase of RF distances for all four approaches, and it affects AAF more than alignment (Figure 1). When the random dropout rate was high, the number of loci that were shared by all samples decreased drastically (Figure S2), resulting in fewer reads being selected (Figure S1). This led to so little data remaining after selection that SBA reads selection performed worse than AAF without selection (Figure 3, *r* = 0.05).

### Real RADseq dataset (Quercus)

For the real RADseq dataset, the phylogenetic relationships between different *Quercus* species given by AAF without reads selection matches fairly closely with the phylogeny constructed using a traditional assembly and alignment method (described in Cavender-Bares et al. 2015) (Figure 5a). Most of the individuals from the same species were grouped together by AAF except some individuals in *Q. geminate* and *Q minima* (Figure 5b). With SBA reads selection (Figure 5c), the normalized RF distance between the AAF and published phylogeny (Figure 5a) decreased from 0.52 to 0.23. Also, in the SBA reads selection reconstruction, individuals from the same species formed their own clades, and the relationship between species was the same as the published phylogeny, except *Q. geminate* and *Q minima* did not form monophyletic clades. This might be due to the introgression that has been reported involving these two species (Cavender-Bares *et al*. 2015). The phylogeny generated by pairwise reads selection is similar to that from SBA reads selection (Figure 5d), with a slightly higher normalized RF distance of 0.29 from the published phylogeny. The lizards dataset also yielded similar phylogeny with original publication (Figure S3).

**Figure 5.**
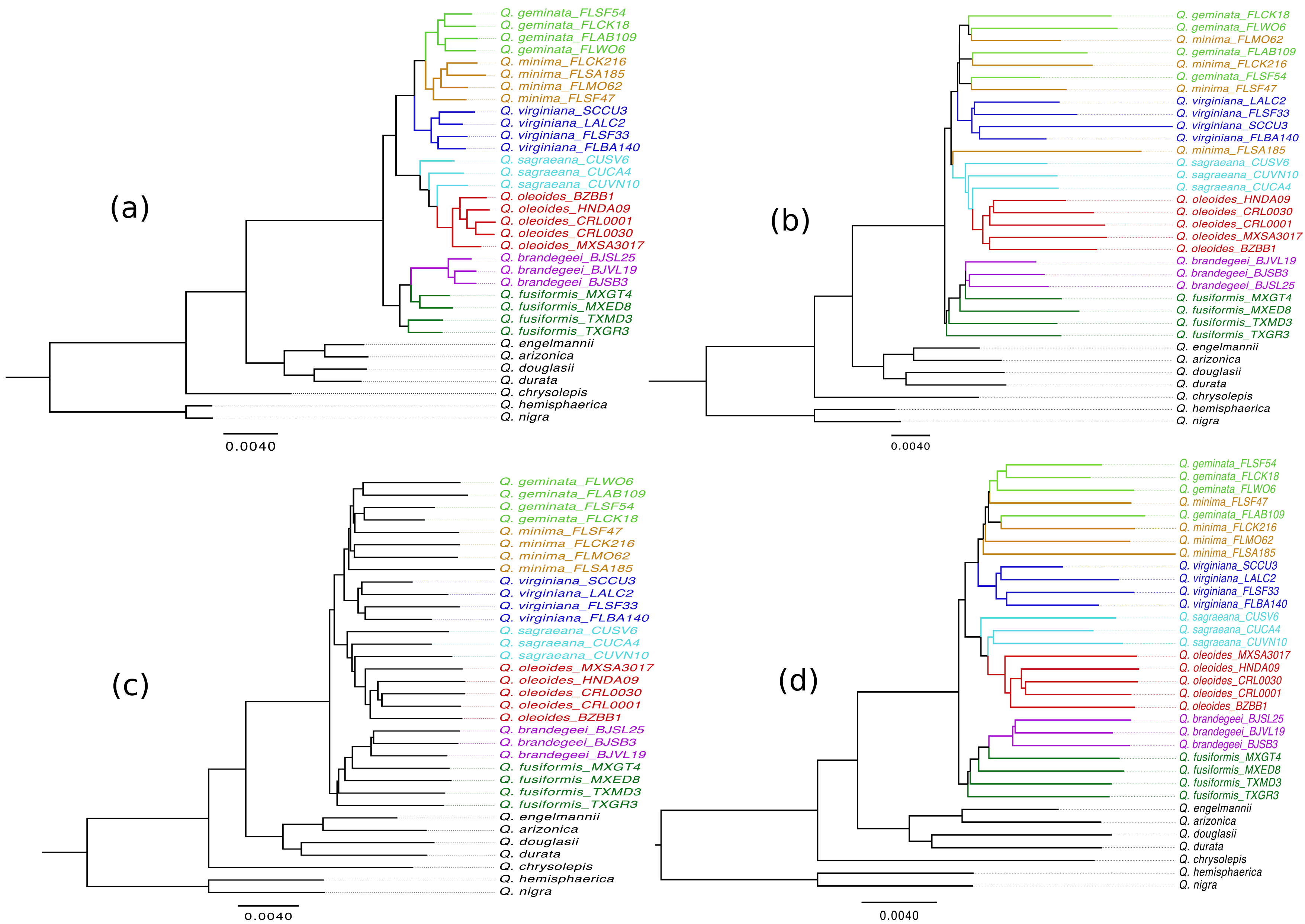
Phylogenies reconstructed using RADseq data from individuals of *Quercus.* (a) Reproduced from the newick from of Figure 2a in Cavender-Bares et al. (2015). (b) Phylogeny reconstructed from the *Quercus* RADseq dataset using AAF without selection (k = 21). (c) Phylogeny reconstructed using AAF with SBA reads selection *(k_s_ = k_p_ =* 21). (d) Phylogeny reconstructed using AAF with pairwise reads selection *(k_s_ = k_p_ =* 21). Individuals from the same species are coded with the same color.

## Discussion

AAF, an assembly and alignment-free method for phylogenetic reconstruction using whole genome data (Fan et al. 2015), was an effective method for analyzing RADseq data even though less data are available to infer phylogenies than with whole-genome sequencing. AAF was further improved for RADseq data by performing reads selection in which reads sharing at least one *k*-mer were assumed to represent the same RAD locus.

This is the first study to investigate the effect of random dropout in addition to nonrandom dropout using simulations. Nonrandom dropouts are caused by mutations that disrupt or generate restriction sites and hence contain phylogenetic information. In contrast, random dropouts occur during the sequencing and data processing stages, and hence contain no phylogenetic information. In other methods that use RADseq data, there is debate about whether to select RAD loci that are shared by a minimum number of samples (Huang & Knowles 2016). This involves a trade-off between quality of data (which increases with selection) and quantity of data (which decreases with selection). Random dropout lowers the quality of the data by adding non-informative noise, and our simulations showed that random dropout increases the advantage of reads selection. While for small samples, sufficient information remains after selecting reads shared by all sample (SBA reads selection), for large samples it is better to select reads between pairs of samples (pairwise reads selection) which retains information regardless of the total sample size. Pairwise reads selection largely overcomes the quality vs. quantity tradeoff, although with the cost of increased computational burden (Table S3).

### AAF vs. alignment

We compared AAF with an alignment-based method for phylogenetic reconstruction using simulations. Because we used simulated data, the alignments we generated were unambiguous, and all included loci were homologous between the different samples. This would not be the case with real RADseq data from non-model organisms because in the absence of a reference genome, the RAD loci have to be identified *de novo*. This can lead to errors when paralogues and repetitive sequences are present. Because we could identify the source of reads in our simulations, the alignment method we used to compare with AAF had an artificial advantage. Nonetheless, we still found that AAF performed slightly better than the alignment-based method when the random dropout rate was low. This could due to two things. First, when there is a heterozygous position in the data, the consensus sequence will contain ambiguous bases and will be ignored in the phylogenetic reconstruction using the alignment-based method. This will result in a loss of information in contrast to the AAF method. Second, in the case of low coverage at a given locus (coverage <= 3), sequencing errors present in a single read would either be kept or make that base pair ambiguous in the consensus sequence, while it would be removed by the filtering step in AAF.

Alignment became more accurate than AAF when the random dropout was high (5% in our simulations). This could be due to the way the two methods treat missing data. While in the alignment method missing data are just ignored, AAF takes into account the presence/absence information in the calculation of evolutionary distance between samples. This is an advantage when a locus is missing due to a mutation in a restriction site (real phylogenetic information), but becomes a drawback when the absence of a locus is due to a low coverage or a random dropout (random noise). When the random noise becomes too large (5% random dropouts), the accuracy of AAF is compromised, and reads selection should be used to remove the noise introduced by missing data.

### Reads selection

Reads selection does not distinguish between random and nonrandom dropout; it keeps loci that are shared by all (Figure 2) or two (pairwise) samples. Therefore, it abandons the phylogenetic information that could be available from nonrandom dropout of loci. Nonetheless, by increasing the proportion of loci that are shared between samples, reads selection improved the performance of AAF in all scenarios in figure 1. This suggests that the impact of including random noise is more detrimental than the loss of real phylogenetic information by excluding missing data.

We tested two methods for reads selection. SBA reads selection is based on the k-mers shared by all the samples in the study. When *k* is sufficiently large to select reads from the same restriction sites (Figure 2), this method leads to substantial improvements in AAF. While this approach is fast and efficient, the number of reads kept when analyzing a large number of samples and/or very divergent species might be too low for accurate phylogenetic reconstruction. In this situation, we recommend the use of pairwise reads-selection that retains more information and hence gives more accurate reconstructions (Figure 3). The drawback of this method is that the computation time scales up quickly as the number of samples increase (Table S2). Furthermore, different loci are used for different pairs of samples, leading to potentially more heterogeneous information when constructing the phylogeny; this could explain why SBA works better than pairwise reads selection when the number of samples is small.

### *k_s_* for reads selection and *k_p_* for reconstruction

The choice of an optimal *k_p_* for phylogeny reconstruction is a balance between reducing homoplasy (favoring larger *k_p_*) and losing information by having multiple evolutionary events on the same *k*-mer (favoring smaller *k_p_*). However, homoplasy is the more severe problem, and using *k_p_* that is too small led to very poor performance (compare *k_p_* = 9 vs. 11 in figure 4 with 20,000 loci). As long as there is enough information, a longer *k_p_* will guarantee little or no homoplasy. This was discussed extensively in Fan et al. (2015).

The choice of an optimal *k_s_* used for selection involves the same balance, although the cost of using a shorter-than-optimal *k_s_* is much less. If *k_s_* is less than optimal, there is little discrimination in the selection of reads, resulting in the same phylogenies as AAF without selection. Once the homoplasy problem is overcome, SBA loci are almost exclusively selected (Figure 2, *k_s_* =15 and 23). Although not all SBA loci were selected, probably due to multiple mutations within *k* (Figure S2, *k_s_* = 15, 23 or 31), there was enough information in the selected SBA loci for accurate phylogeny reconstructions.

### Appropriate scale for AAF

RADseq was originally designed for population genetics. In recent years, it has been used in at least a dozen interspecific phylogeny reconstruction studies including both plants (Eaton & Ree 2013; Hipp *et al*. 2014) and animals (Ebel *et al*. 2015), usually at the genus level. Initially, it was reported that RADseq data were not appropriate for reconstructing deep phylogenies (Rubin *et al*. 2012). The concern for distantly related species is that there might not be enough restriction sites shared among species due to accumulated mutations in restriction sites. However, when evolutionary divergence is small, even though restriction sites are retained, there might not be enough informative molecular variation to infer evolutionary history. Therefore, there has been no clear consensus on what depths of divergences are appropriate for RADseq analyses.

In our simulations we used two ultrametric trees with a tip-to root depth of 0.1 and 0.01, representing deep (comparable to the divergence times of 100mya for primates, Fan et al. 2015) and shallow (intraspecific) evolutionary distances. The real RADseq data we used for demonstration, the *Quercus* dataset from Cavender-Bares et al. (2015), has an average tip-to-root distance of 0.028 mutations per site, between the two levels in our simulations. In contrast to concerns about RADseq being only appropriate for shallow divergences, our simulated RADseq data allowed accurate reconstruction with deep phylogenies, and fewer loci were required (Figure 1). Previous studies suggested that using RADseq for deep phylogenies might be limited due to the challenge of de-novo assembling the flanking regions around the restriction sites from short sequences that are too divergent, and the large amount of missing data due to mutations in the restriction sites. Using AAF there is no need for assembly, and reads selection helps with the problem with missing data. Therefore, AAF is suited for phylogenetic analysis of RADseq data with deeper divergence.

At the other end of the divergence range, shallow phylogenies are difficult to reconstruct due to the small amount of phylogenetic information accumulated over short evolutionary scales. As a consequence, the accuracy of the reconstructed phylogenies is lower for shallow divergences (Figure 1). Although AAF outperformed the alignment method we used, neither method provided satisfactory results. More loci are required to accurately reconstruct shallower phylogenies. This could be achieved by using either more restriction enzymes or enzymes with more frequent recognition sequences.

## Conclusion

In both simulated and real datasets, AAF produced accurate reconstructions of phylogenetic relationships between samples using RADseq data, especially when aided by reads selection. Computationally, AAF is much computationally more efficient than traditional alignment-based methods. Furthermore, AAF with reads selection requires few decisions about parameter choice, which is confined to selecting *k_s_* and *k_p_* using simple rules; this guards possible user bias. Finally, the approaches we have presented should work well on other kinds of reduced-representation genomic data.

## Data accessibility

Tree files from figure 5 and figure S3 can be found in the Dryad data repository at http://datadryad.org (doi:10.5061/dryad.r0hq0).

## Supporting information

Additional supporting information may be found in the online version of this article.

---

**Figure S1.**
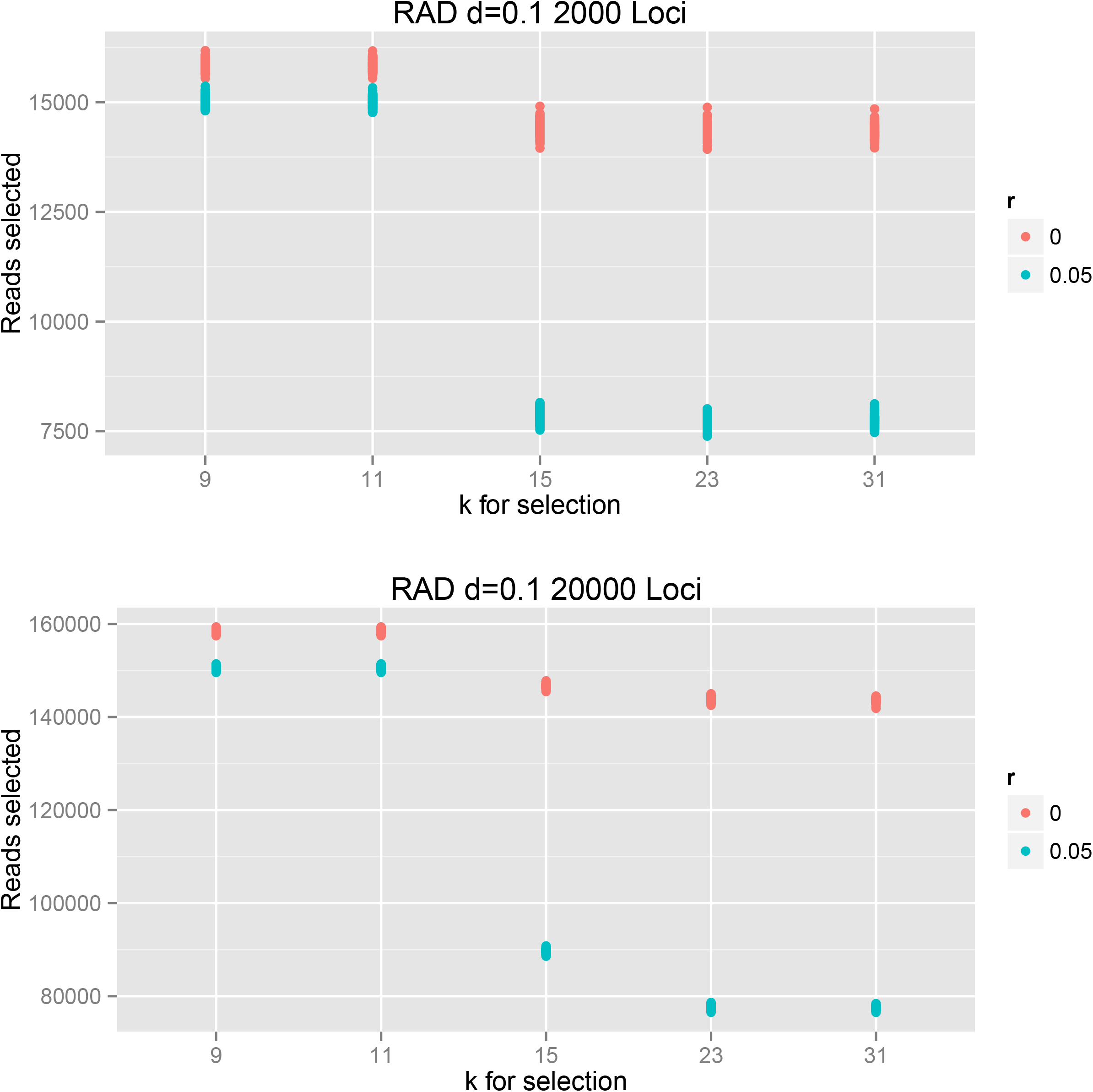
Number of reads selected with different *k_s_*. The simulations are based on the deep 12-species starting tree (*d* = 0.1).

**Figure S2.**
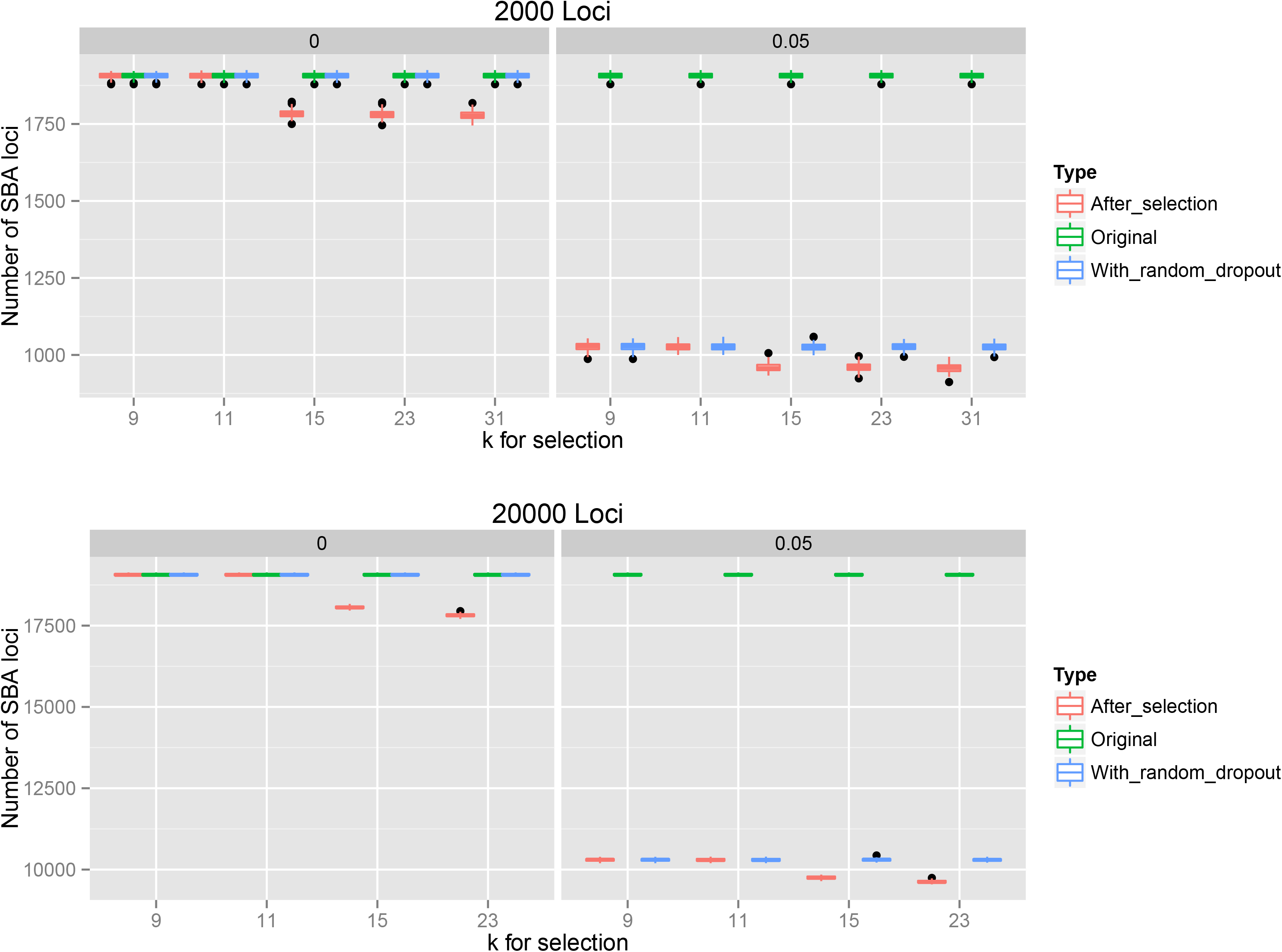
Number of SBA loci in the original data (SBA_b), after adding random dropout rate (SBA_r), and after selection (SBA_s) with different *k_s_*. The simulations are based on the deep 12-species starting tree (*d* = 0.1).

**Figure S3.**
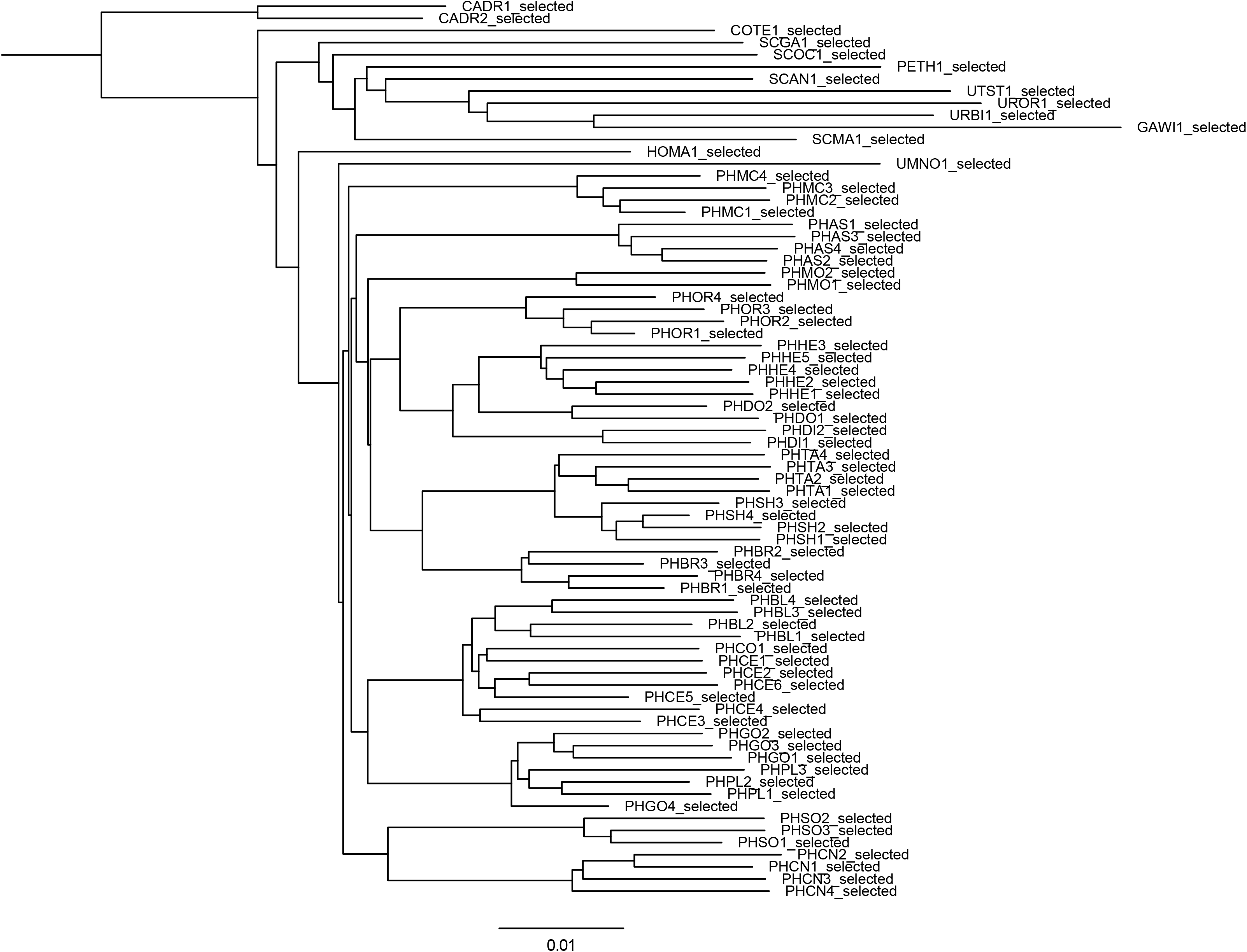
Phylogenies reconstructed using RADseq data from Lizards.

**Table S1** Metadata of the *Quercus* dataset from Cavender-Bares *et al*. (2015)

**Table S2** Metadata of the Lizard dataset from Leache *et al*. (2015)

**Table S3** Mean memory and computational time required by each method using one core.

